# Genetic Characterization and Curation of Diploid A-Genome Wheat Species

**DOI:** 10.1101/2021.08.20.457122

**Authors:** Laxman Adhikari, John Raupp, Shuangye Wu, Duane Wilson, Byron Evers, Dal-Hoe Koo, Narinder Singh, Bernd Friebe, Jesse Poland

## Abstract

The A-genome diploid wheats represent the earliest domesticated and cultivated wheat species in the Fertile Crescent and the donor of the wheat A sub-genome. The A-genome species encompass the cultivated einkorn (*Triticum*. *monococcum* L. subsp. *monococcum*), wild einkorn (*T. monococcum* L. subsp. *aegilopoides* (Link) Thell.) and *T. urartu*. We evaluated the collection of 930 accessions in the Wheat Genetics Resource Center (WGRC), using genotyping-by-sequencing (GBS) and identified 13,089 curated SNPs. Genomic analysis detected misclassified and duplicated accessions with most duplicates originated from the same or a nearby locations. About 56% (n = 520) of the WGRC A-genome species collections were duplicates supporting the need for genomic characterization for effective curation and maintenance of these collections. Population structure analysis confirmed the morphology-based classifications of the accessions and reflected the species geographic distributions. We also showed that the *T. urartu* as the closest A-genome diploid to wheat through phylogenetic analysis. Population analysis within the wild einkorn group showed three genetically distinct clusters, which corresponded with wild einkorn races α, β, and γ described previously. The *T. monococcum* genome-wide F_ST_ scan identified candidate genomic regions harboring domestication selection signature (*Btr1*) on the short arm of chromosome 3A^m^ at ~ 70 Mb. We established A-genome core set (79 accessions) based on allelic diversity, geographical distribution, and available phenotypic data. The individual species core set maintained at least 80% of allelic variants in the A-genome collection and constitute a valuable genetic resource to improve wheat and domesticated einkorn in breeding programs.

**One-sentence summary:** Genotyping of gene bank collections of diploid A-genome relatives of wheat uncovered relatively higher genetic diversity and unique evolutionary relationships which gives insight to the effective use of these germplasm for wheat improvement.

## Introduction

Wheat wild relatives are an important reservoir of genetic diversity that can be utilized for wheat improvement, particularly for diseases, insect pests, and abiotic stress tolerance (Wulff & Moscou, 2014). Cultivated tetraploid (pasta wheat, *Triticum turgidum*) and hexaploid (bread wheat, *Triticum aestivum*) wheat arose through successive whole-genome hybridization between related species in the *Triticeae*. Although polyploidization in wheat enabled broad adaptation and genome plasticity found in polyploids (Comai, 2005), it also created severe genetic bottlenecks within each subgenome (Feldman & Levy, 2012). Likewise, of the three natural races within wild einkorn, only one natural race (β) has been domesticated, thus, genetic diversity in the wild einkorn is expected to be greater than in domesticated einkorn (Pourkheirandish et al., 2018). Some recent findings, however, reported no or low reduction in nucleotide diversity through einkorn domestication, most likely indicating a minimal bottleneck during domestication of cultivated einkorn (Kilian et al., 2007). This was true when diversity comparisons were performed between wild einkorn specific races (α and β) vs. domesticated einkorn. However, when the comparison was made between the domesticated einkorn vs. all groups of wild einkorn, the wild einkorn diversity was much higher than found in the cultivated accessions. The value of A-genome species diversity for alleviating the wheat diversity bottleneck have been described (Brunazzi et al., 2018; Mondal et al., 2016). Thus, diversity assessment in germplasm collections of diploid A-genome species is crucial for conservation planning and efficient utilization of germplasm in breeding.

A-genome wheat species (2n = 2x = 14, AA) are diploid grasses including the wild einkorn (*T*. *monococcum* L. subsp. *aegilopoides* (Link) Thell.), domesticated einkorn (*T*. *monococcum* L. subsp. *monococcum*), and *T*. *urartu* (van Slageren, 1994). Molecular and cytological studies have confirmed that *T. urartu*, a related species sharing the same genome as domesticated einkorn, is the A-genome ancestor to cultivated wheat (*T*. *aestivum*) (Dong et al., 2012). In the first polyploidization event that occurred ~500,000-150,000 million years ago (MYA) (Charmet, 2011), *T*. *urartu* naturally hybridized with a B-genome donor grass, an extant species but close relative of *Aegilop*s *speltoides* Tausch, giving rise to the wild tetraploid wheat *T*. *turgidum*L. subsp. *dicoccoides* (Körn. Ex Asch. & Graebn.) Thell. (AABB, 2n = 4x = 28) (Nair, 2019). In the next event, the cultivated tetraploid emmer wheat (*T*. *turgidum* subsp. *durum* (Desf.) Husn.) naturally hybridized with the D-genome donor species (*Ae*. *Tauschii* Coss) forming hexaploid bread wheat (AABBDD, 2n = 6x = 42). The A-genome species morphologically resemble cultivated tetraploid and hexaploid wheat more than any other surviving diploid genome donors and are predominant in the Fertile Crescent (Heun et al., 1997). Domestication of einkorn wheat, together with emmer wheat and barley around 12,000 years ago, transformed human culture from hunting-gathering to agriculture, popularly known as the ‘Neolithic Revolution’ (Kilian et al., 2010). The Karacadağ mountain in the southeast Turkey has been considered the geographical point for einkorn domestication (Brandolini et al., 2016).

The donor of the A genome of the bread wheat, *T*. *urartu*, is estimated to have diverged nearly 0.57 – 0.76 MYA from another widespread A-genome diploid species, *T*. *monococcum*. Interspecific crosses between *T. urartu* and *T. monococcum* are infertile, confirming the large phylogenetic distance and genetic differentiation of the species (Middleton et al., 2014). Like hexaploid wheat, A-genome species have a large genome size with a mean nuclear DNA content of 5.784 pg/1C in *T*. *urartu* to 6.247 pg/1C in *T*. *monococcum* subps. *aegilopoides* (Özkan et al., 2010). Morphologically, *T*. *urartu* possesses smooth leaves, a brittle rachis, and smaller anthers (< 0.3 mm). The wild einkorn (*T*. *monococcum* subsp. *aegilopoides*) are characterized with a brittle rachis, hairy leaves, and larger (≥0.5 mm) anthers. Domesticated einkorn has a nonbrittle (semitough) rachis with smooth leaves (Brandolini & Heun, 2019).

Being homologous to the wheat A-genome, these species provide useful sources for wheat improvement using wide crosses and cytogenetics approaches. The A-genome species are important genetic resources for pest resistance and stress tolerance. For example, *T*. *urartu* was identified as a source of resistance to the root lesion nematode *Pratylenchus thornei* (Sheedy et al., 2012) and stem rust (Rouse & Jin, 2011). Novel stem rust resistance genes *SrTm5* and *Sr60* were mapped in an F_2_ population derived from crosses between wild and the cultivated einkorn (Chen et al., 2018). Sr35, the first gene cloned against the devastating stem rust race UG99, also originates from *T*. *monococcum* (Saintenac et al., 2013). A leaf rust gene, *Lr63*, in wheat chromosome 3AS was introgressed from *T*. *monococcum* (Kolmer et al., 2010). Surveying the genetic variation in A-genome species that can be utilized in wheat improvement has lagged, considering the potential value of more effectively utilizing these species for wheat improvement.

Einkorn has multiple botanical names in the literature as proposed by the various taxonomists, and confusion related to the einkorn nomenclature is widespread. In 1948, Schiemann classified einkorn as wild einkorn (*T*. *boeoticum* subsp. *thaoudar*), the feral einkorn (*T*. *boeoticum* subsp. *aegilopoides*), and the domesticated einkorn (*T*. *monococcum* subsp. *monococcum*) (Brandolini et al., 2016; Schiemann, 1948). MacKey published einkorn classification in 1954 (Key, 1954) and updated the nomenclature several times through 2005 (Mac Key, 2005a). van Slageren also published the einkorn nomenclature, where the wild and domesticated einkorn were simply named as *T*. *monococcum* L. subsp. *aegilopoides* (hereafter subsp. *aegilopoides*) and *T*. *monococcum* L. subsp. *monococcum* (hereafter subsp. *monococcum*), respectively (van Slageren, 1994). In this study, we follow van Slageren’s (1994) einkorn taxonomy, because the A-genome species collection in the Wheat Genetics Resource Center (WGRC) at Kansas State University (KSU) were initially classified using this nomenclature (van Slageren, 1994).

A well-characterized population structure of A-genome species is critical to formulating effective conservation strategy, selecting diverse germplasm, and enhancing the accuracy of the genomic analysis with structure information (Singh et al., 2019). Population structure and diversity assessment have become easier with next-generation sequencing, which makes discovery of thousands of genotyping markers possible. Here, we used genotyping by sequencing (GBS) for single nucleotide polymorphism (SNP) discovery. GBS is straightforward, high-throughput, and with multiple downstream pipelines for data processing (Poland et al., 2012a). However, population structure of A-genome species has not been evaluated in detail with the resource of whole-genome profiling. Therefore, our objectives are to : i) curate A-genome wheat accessions in the gene bank by identifying duplicates and misclassified accessions, ii) assess the population structure and genetic diversity of the A-genome wheat species, and iii) establish genetically, geographically, and phenotypically representative core collections for A-genome species within the WGRC gene bank.

## Results

### A-Genome Species Distribution

Most of the wild einkorn (subsp. *aegilopoides*) in our collection, were collected across Turkey, northern Iraq, west Iran, and Transcaucasia, whereas the majority of domesticated einkorn (subsp. *monococcum*) were from west Turkey and the Balkans (Figure 1, Supplementary Table S1). About half of the *T*. *urartu* accessions were from eastern Lebanon, around the Beqaa Valley, and a major part were from southeast Turkey (Figure 1). The A-genome species are known to span from Transcaucasia through Anatolia to the Caspian Sea. The WGRC collection covers the geographic range of this species. After genomic characterization including misclassified accessions adjustment, we retained 196 *T*. *urartu*, 145 of domesticated einkorn, and 584 wild einkorn (Supplementary Table S1). There were also 5 tetraploids identified in the population which were curated to correct species.

**Figure 1.**
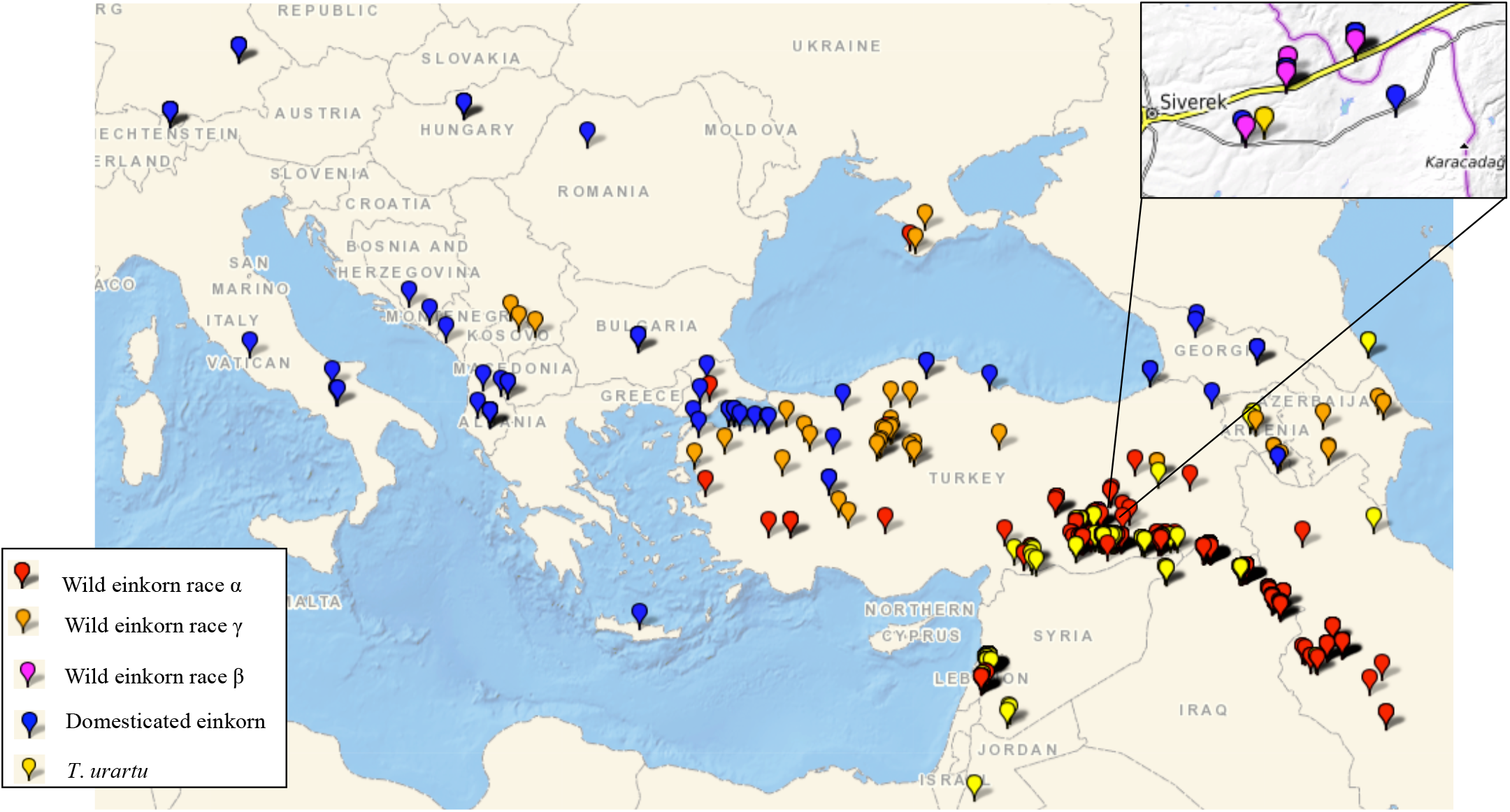
Geographic distribution of A-genome wheat species in the WGRC gene bank. Collection sites of accessions in this study are designated for domesticated einkorn (*Triticum*. *monococcum* subsp. *monococcum*) (red); α race within wild einkorn (*T*. *monococcum*. subsp. *aegilopoides*) (blue); γ race wild einkorn (orange); β race wild einkorn (magenta); and *T*. *urartu* (yellow).

### Markers and Genotyping

For all A-genome accessions, we identified 44,215 biallelic SNPs after a filter for passing Fisher exact test of disassociated alleles. Separating this by subspecies, we had 24,314 biallelic SNPs for subsp. *aegilopoides*, 19,940 biallelic SNPs for *T*. *urartu*, and 13,957 biallelic SNPs for subsp. *monococcum*. Upon filtration (MAF > 0.01, 30% < missing, 10% < heterozygosity), we retained 7432 SNPs for *T*. *urartu* and 6734 SNPs for *T*. *monococcum*, 6343 SNPs for subsp. *aegilopoides*, and 3980 for subsp. *monococcum*. For wheat and A-genome diploids together we found 15,300 filtered SNPs. For A-genome species diversity assessment, thousands of segregating loci were available for the groups defined by population analysis and core set selections (Table 1). We filtered the loci for MAF (MAF > 0.01) before splitting the VCF file to the species and sub-species and observed loci that were fixed or otherwise one heterozygous genotype call within the individual species and subspecies. To compute total segregating loci per group and minimize the effect of potential sequencing error, we did further filtration and removed any loci that were segregating only due to a single heterozygous genotype and otherwise the major allele is fixed in remaining population (Table 1).

**Table 1.** A-genome species and sub-species groups with number of samples, the Nei’s diversity indices, and number of segregating loci. The percentage of segregating SNPs for core set groups were estimated relative to the segregating loci within the respective groups.

| Group | Number of<br>Samples | Diversity<br>Index | Segregating<br>SNPs |
| --- | --- | --- | --- |
| **A-genome species (*T. monococcum* + *T. urartu*)** | **925** | **0.25** | **13089** |
| *T. monococcum* (einkorn) | 729 | 0.106 | 6587 (50.3%) |
| Domesticated einkorn (subsp. *monococcum*) | 145 | 0.058 | 3637 (27.8%) |
| Wild einkorn (subsp. *aegilopoides*) | 584 | 0.086 | 6213 (47.4%) |
| α race einkorn | 524 | 0.073 | 5119 (39.1%) |
| γ race einkorn | 48 | 0.093 | 4440 (33.9%) |
| β race einkorn | 12 | 0.058 | 2622 (20.1%) |
| *T. urartu* | 196 | 0.069 | 4072 (31.11%) |
| **A-genome species core set** | **79** | **0.271** | **12907 (98.6%)** |
| *T. monococcum* core | 60 | 0.116 | 5926 (89.9%) |
| Wild einkorn core | 41 | 0.098 | 5324 (85.6%) |
| Domesticated einkorn core | 19 | 0.065 | 3039 (83.6%) |
| *T. urartu* core | 19 | 0.072 | 3324 (81.6%) |

### Gene Bank Curation

We identified and corrected a total of 22 misclassified accessions using fastStructure analysis, phylogenetic and PCA clustering (Supplementary Figure S1) including nine *T*. *urartu*, two subsp. *monococcum*, six subsp. *aegilopoides*, and five tetraploid accessions (Supplementary Table S3). As large number of accessions in both *T*. *urartu* and subsp. *aegilopoides* collection were from southeast Turkey; we observed most of the misclassified accessions also were from the same site.

While evaluating the collection for duplicate accessions, we compared various number of loci for allele matching per A-genome species (Table 3) as the SNPs were filtered to keep only the sites with > 0.05 MAF, < 50% missing and < 10% heterozygous. We identified and used a threshold of ≥ 99% identity by state (IBS) to declare the individuals as identical accessions to warrant the inclusion of identical accessions in the duplicate set (Supplementary Figure S2) with tolerance for sequencing and genotyping error. With these criteria we identified a total of 520 (56%) duplicated accessions which were mostly observed within *T*. *urartu* and within α race subsp. *aegilopoides* (Supplementary Table S1). To confirm this analysis, we checked the collection sites of the groups of duplicates identified and all of the respective sets of duplicates were collected from the same or nearby sites. We further observed the duplicates had same phenotypes as the glume color scores were the same for sets of duplicates (Supplementary Table S1), confirming the accuracy of using the GBS data for identification of duplicated accessions. For instance, TA471 and its 11 duplicates had glume color score of 7 while on a scale of 1 (white) to 9 (black) (Supplementary Table S1).

### Relationship Between A-genome Diploid and Wheat

The genetic grouping of A-genome diploids and CIMMYT wheat lines together showed that wheat is closer to *T*. *urartu* than to *T*. *monococcum* (Supplementary Figure S3), a finding in agreement with the known relationship between the species. The unrooted NJ tree constructed for wheat and A-genome diploid wheat showed five accessions (TA282, TA10915, TA1325, TA1369, and TA10881) clustering far from the *T*. *urartu* major clade (Supplementary Figure S3). Cytological analysis identified them as tetraploid (2n=28) (Supplementary Figure S4). Therefore, we excluded these five accessions from population analysis. This observation confirms that GBS also enables identifying cryptic accessions with different ploidy levels in the population.

### A-genome Population Structure and Wild Einkorn Genetic Races

Population grouping in the fastStructure analysis at K=2 to K=7 showed the A-genome genetic structure was split with the known biological and geographical characterization (Figure 2). This analysis revealed a number of misclassified accessions that were individually curated and checked, including morphological confirmation, and were reclassified to the appropriate group (Supplementary Table S3).

**Figure 2.**
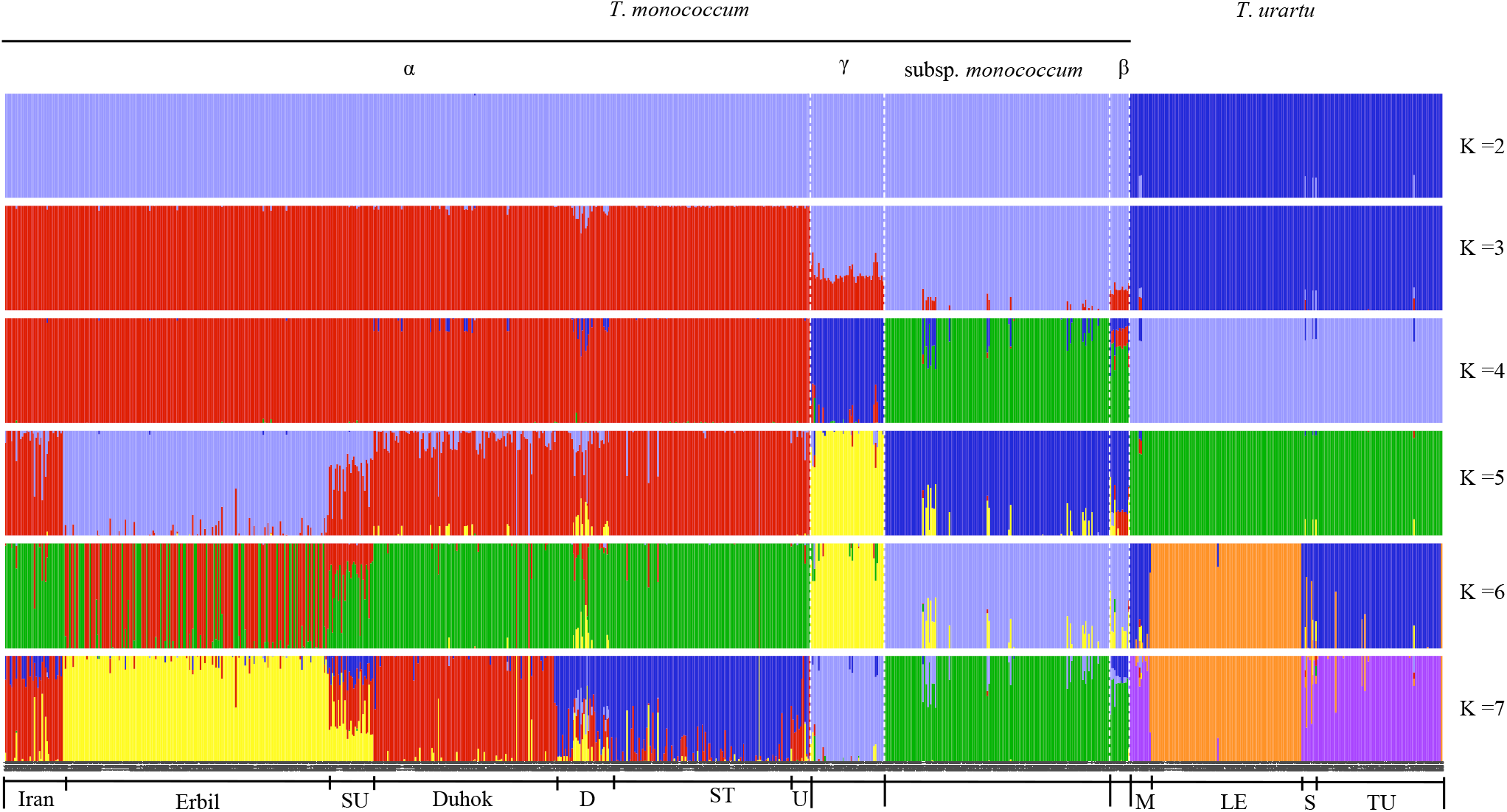
Population structure of A-genome wheat species: *Triticum monococcum* L. and *T*. *urartu*. Subpopulations were determined using fastStructure at K=2 to K=7. Each color represents a population, and each bar indicates the admixture proportion of an individual accession from K populations. The subgroup within α, which is exemplified by yellow color sole includes the accessions from Erbil (also spelled Arbil), Iraq, whereas the subgroup embodied by red color only comprises the accessions from Duhok (ancient name ‘Dahuk’, Iraq). The bars with purple color only represent the accessions from southeast Turkey (ST). Other admixture types within α included accessions were from Iran, SU (Sulaymaniyah (Iraq)), random different sites (D) and unknown sites (U) as indicated. Within *T*. *urartu*, the LE group represents accessions from Lebanon, the TU includes accession from Turkey, S indicates accessions from Syria and M shows accessions from mixed sites.

At K=2, the population differentiation occurred only at the level of species, the accessions split into *T*. *monococcum* and *T*. *urartu*, confirming known species differences (Figure 2). At K=3, the two subspecies of *T*. *monococcum* differentiated with the accessions in the α wild einkorn race were clearly differentiated from domesticated einkorn. However, the other races of wild einkorn (β and γ) appeared to be an admixture, supporting that there is not complete differentiation between the wild and domesticated einkorn, a classification that is simply based on the few morphological characteristics of the domestication syndrome.

We observed differentiation of wild einkorn into genetically distinct groups at K=7. Comparing these three wild einkorn subgroups with the α, β, and γ wild einkorn races described by (Kilian et al., 2007), we report the three genetic subgroups as representing the races α, β, and γ by identifying common USDA Plant Introduction (PI) numbers for accessions in both studies. The genetic clustering pattern and geographical distribution then confirmed that the subgroups within subsp. *aegilopoides* represents α, β, and γ races described and we hereby name these genetic groups accordingly (Supplementary Table S4) (Kilian et al., 2007). In (Kilian et al., 2007), the α race accessions were primarily from southeast Turkey, northern Iraq, and Iran; the γ race involves accessions from Transcaucasia to western Anatolia; and the β race comprises a few accessions collected around Karacadag Turkey (Figure 1, Supplementary Table S1). Based on population differentiation, α race exhibited the strongest differentiation with domesticated einkorn and should represent the base population of subsp. *aegilopoides*, whereas the β race of wild einkorn exhibited the least differentiation with subsp. *monococcum*. Interestingly, the β race did not fully differentiate from subsp. *monococcum* at any value of K (Figure 2), supporting that domesticated einkorn originated out of this subpopulation, which already largely differentiated from the other wild einkorn, or (2) that the β race represents ‘feral’ subsp. *monococcum* accessions that were, at one point, fully domesticated but reverted to wild plant types through introgression and admixture.

At K=5, the population subgrouping according to the accession origin was observed in α race accessions within the wild einkorn. Accessions from Erbil (ancient name ‘Arbil’) differentiated as a subpopulation, and the accessions from Sulaymaniyah (Iraq) split as the admixture of the Erbil subgroup and the remaining accessions at K=5 (Figure 2). We could not observe any new differentiation within the wild einkorn group at K=6. However, at K=7, we observed three distinct subgroups and a higher level of admixture within the α race of subsp. *aegilopoides* (Figure 2). Also, there were two main sets of admixture types; the first set mainly consists of accessions from Iran that shared ancestry from the Duhok (red) and Turkey (purple) subgroups, and the second corresponds with accessions from Sulaymaniyah (Iraq) and has ancestry from all three subgroups. Hence, within the population of α race einkorn accessions, three subgroups exist; Erbil, Duhok, and Turkey, and two groups of genetic admixtures (Iran and Sulaymaniyah), named from their origin.

We did not observe any subgrouping within the accessions from the southeast Turkey, yet the accessions were primarily from two sites (Sanliurfa and Mardin). The grouping pattern of three subgroups within the α race accessions provided a new insight into the wild einkorn subgrouping and their genetic relationships. We did not observe within population differentiation in domesticated einkorn group.

In *T*. *urartu*, the subgrouping occurred at K=6, and was unchanged at K=7 (Figure 2). Two major *T*. *urart<u>u</u>* subgroups represented accessions from Turkey (#T) and another from Lebanon (#L). Few *T*. *urartu* accessions were from Syria (#S); some showed admixture, and some had a clean ancestry that resembled accessions from Turkey (Figure 2). The few remaining accessions primarily were from Transcaucasia (#M) and exhibited an ancestry similar to accessions from Turkey (Figure 2).

### Phylogenetic Clustering and PCA

The phylogenetic clustering split the A-genome accessions into separate clades for *T*. *urartu*, *T. monococcum* subsp. *monococcum*, and all races within the subsp. *aegilopoides* (Figure 3). Only 12 accessions were retained within race β, and the accessions were clustered with some other domesticated einkorn accessions (Figure 3). The *T*. *urartu* clade distantly clustered in both PCA and phylogenetic analysis from either of the einkorn clade indicating the obvious genetic differences between species. The misclassified accessions (Supplementary Figure S1) observed in the phylogenetic clustering were re-classified into proper genotype-based classes.

**Figure 3.**
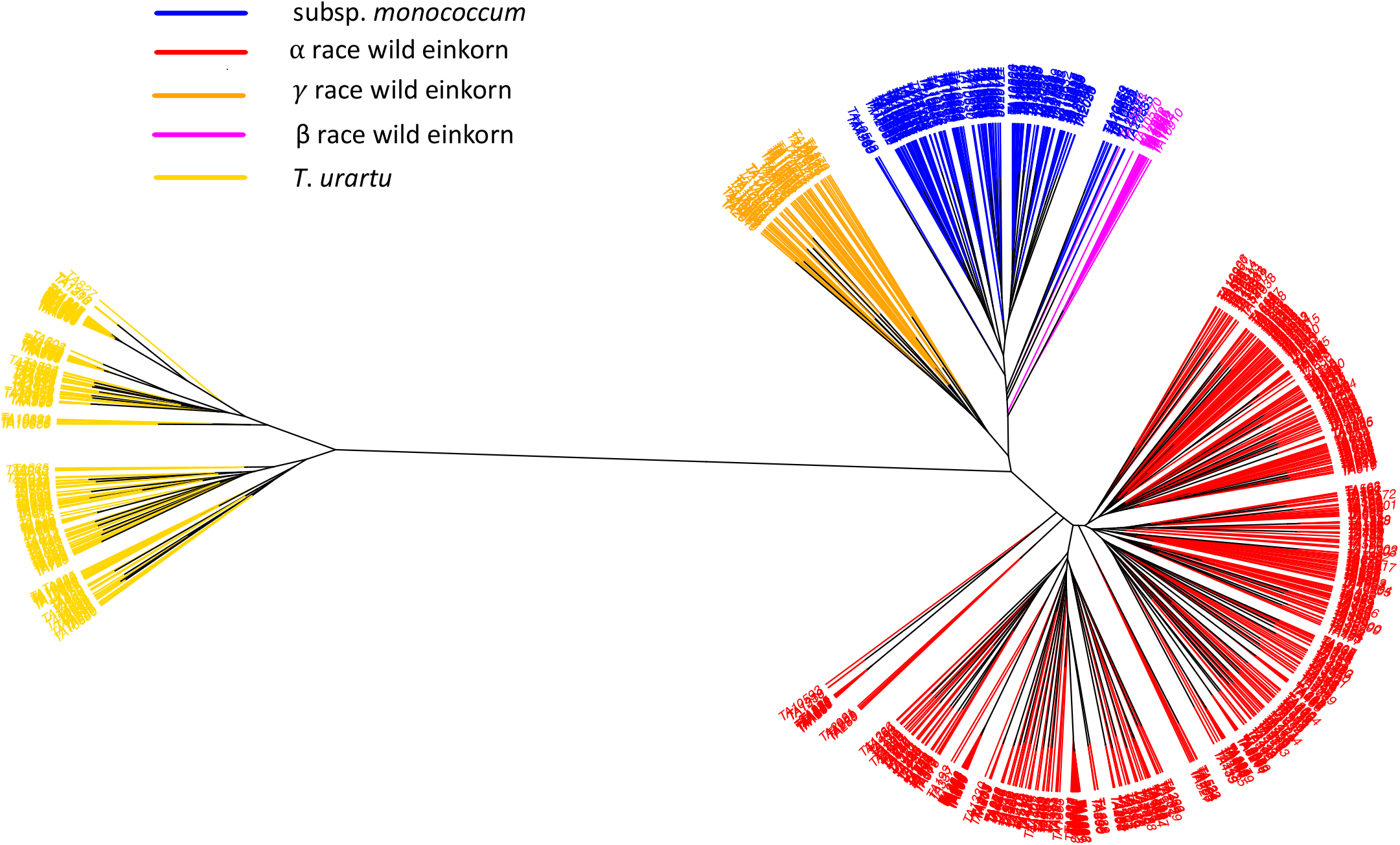
An unrooted Neighbor-Joining (NJ) tree of A-genome species: *T*. *urartu*, subsp. *aegilopoides*, and subsp. *monococcum*. The tree branches are colored based on the genetic grouping of the accessions after correcting misclassified accessions. *T*. *urartu* (yellow), domesticated einkorn (red), wild einkorn race α (blue), and wild einkorn race γ (green) are shown.

**Figure 4.**
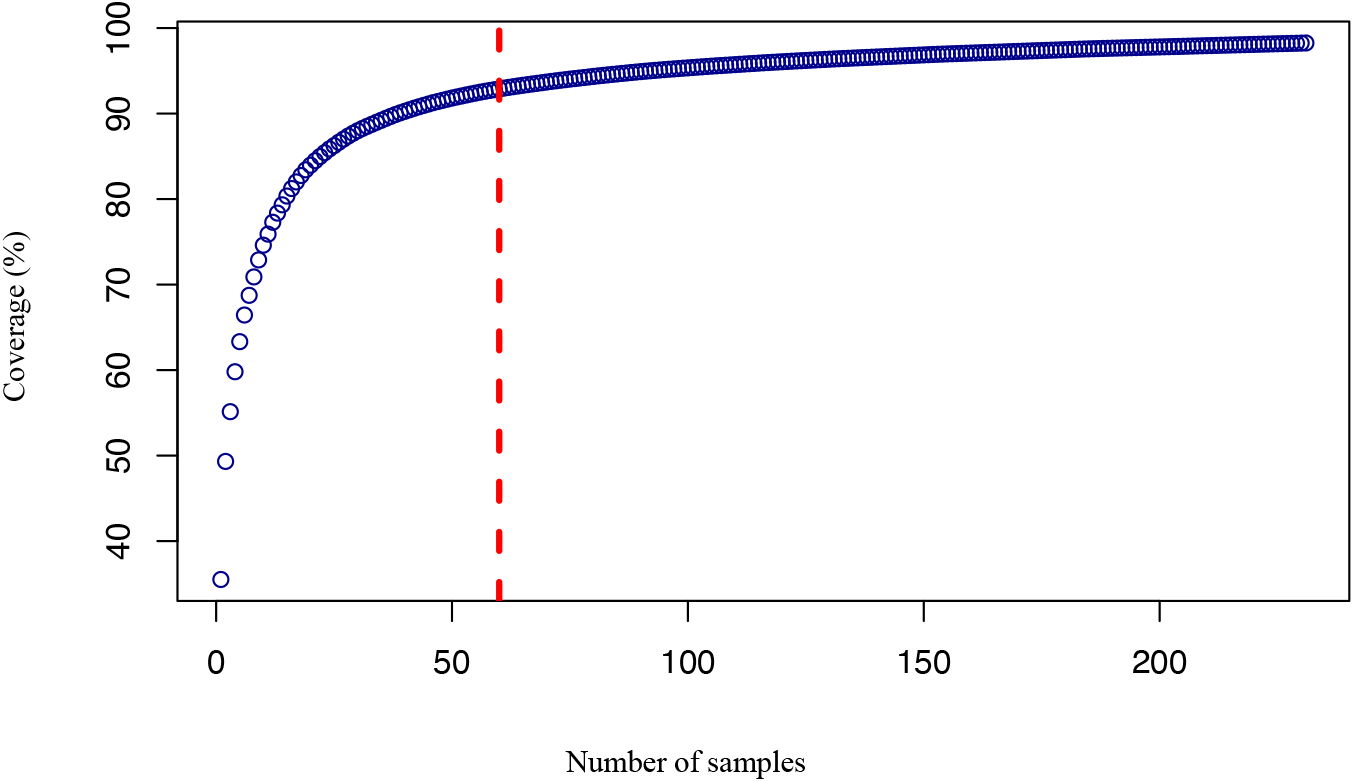
Relationship between the allele coverage as estimated using GenoCore and the number of samples selected in the core for einkorn group (*T*. *monococcum*). The threshold for 60 accessions at approximately 90% genotype coverage is shown with vertical red line.

A PCA plot of A-genome species also showed accessions clustering as in fastStructure and phylogenetic analysis (Supplementary Figure S5). The first principal component (PC1), which grouped the accessions of *T*. *monococcum* and *T*. *urartu* in two primary clusters, explained 58% of the variation. The PC2, which divided the einkorn accessions, explained 8% of the variation and separated domesticated and different races within the wild einkorn. Misclassified accessions previously observed also were revealed in the PCA analysis and their taxonomy classification adjusted.

### Genetic Diversity and F_ST_

A considerably high Nei’s diversity index (0.25) was observed for the complete set of A-genome accessions. The Nei’s diversity indices for individual A-genome species ranged from 0.058 for domesticated einkorn to 0.106 for the entire einkorn group. Among the three races of wild einkorn, the Nei’s diversity indices of β race (0.058) was the lowest and γ was the highest (0.093; Table 1). As expected for diverse accessions, we found a high density of alleles with low minor allele frequency (MAF) (Supplementary Figure S6).

Population differentiation within the A-genome species were further verified by pairwise fixation index (F_ST_) values (Nei’s, 1987) computed between the groups. Pairwise F_ST_ between *T*. *urartu* and entire einkorn were greater than 0.80, supporting that the two species are strongly differentiated (Table 2). The pairwise F_ST_ (0.56) between the α race and domesticated einkorn indicated the strongest differentiation between any two groups within the einkorn, whereas the weakest differentiation (F_ST_ = 0.31) was between the β race and domesticated einkorn, supporting the model that this wild race was the most likely forerunner of domesticated einkorn as previously hypothesized (Kilian et al., 2007). The concept also was endorsed by the origin of β race einkorn in the WGRC collection, mostly from Diyarbakir and Sanliurfa, which are near Karacadag and Kartal-Karacadag mountains (points of domestication). Nonetheless, the genetic grouping of β also occurred with subsp. *monococcum* in the unrooted NJ tree (Figure 3). Pairwise F_ST_ (~ 0.40) between pairs: ‘γ race - subsp. *monococcum*’ and ‘γ race - α race’ implicit the differentiation of γ race as a genetically intermediate type from truly wild α race and domesticated einkorn (Table 2). The pairwise F_ST_ computed between two subpopulations (Turkey and Lebanon) of *T*. *urartu* was 0.52, which also agrees with the population structure analysis.

**Table 2.** Pairwise F_ST_ coefficients among the A-genome wheat species. Higher F_ST_ reflects a stronger population differentiation. The α, β and γ genetic races comprise the wild einkorn (*T*. *monococcum* subsp. *aegilopoides* L.).

| | $\alpha$ race | $\gamma$ race | $\beta$ race | *T. urartu* |
| --- | --- | --- | --- | --- |
| subsp. *monococcum* | 0.56 | 0.41 | 0.31 | 0.87 |
| $\alpha$ race | - | 0.40 | 0.50 | 0.86 |
| $\gamma$ race | - | - | 0.37 | 0.83 |
| $\beta$ race | - | - | - | 0.86 |

**Table 3.** Number (#) of unique accessions, number of accessions in a duplicate set consisting maximum identical accessions, and total accessions of A-genome species: *T*. *urartu*, domesticated einkorn (subsp. *monococcum*), and wild einkorn (subsp. *aegilopoides*) three genetic races: α, γ, and β. The identical accessions were detected using pairwise allele matching.

| | $\alpha$ race | $\gamma$ race | $\beta$ race | subsp. *monococcum* | *T. urartu* |
| --- | --- | --- | --- | --- | --- |
| **Total accessions** | **524** | **48** | **12** | **145** | **196** |
| # Loci compared | 4112 | 4112 | 4112 | 3337 | 6356 |
| Max duplicates set | 28 | 3 | 0 | 5 | 39 |
| **Unique accessions** | **198 (37.8%)** | **37 (77%)** | **12 (100%)** | **97 (66.8%)** | **61 (31.2%)** |

Pairwise F_ST_ computed between the subpopulations within α race of subsp. *aegilopoides* signaled out the geographical differentiation and the potential gene flow within this wild einkorn race. Consistent with the fastStructure output, the Erbil subgroup showed the stronger differentiation (higher F_ST_) with other wild einkorn subgroups (Supplementary Table S5). The subgroup Duhok and southeast Turkey and their admixture group (Iran) had the minimum pairwise F_ST_ (~ 0.12). The accessions within the admixture group of Sulaymaniyah displayed almost similar differentiation (~F_ST_ = 0.16) with three subgroups, which agrees with population structure as the admixture group has ancestry from all three.

### F_ST_ Scan and Einkorn Selection Signature

After filtration and imputation, we had 6,622 SNPs segregating in *T*. *monococcum* on which we calculated per site F_ST_ values for each of the seven chromosomes that ranged from near 0 to 1. Both methods, Porto-Neto et al. (2013) and VCFtools, produced similar results for raw and smoothed F_ST_ values. We used a genome-wide threshold of 3σ (0.24) over the mean F_ST_, from which we observed only a single-selection signature on short arm of chromosome 3A (Supplementary Figure S7) after smoothing using Lowess method (f = 0.1) (Pintus et al., 2014). This selection signature corresponded to the locus that harbors the brittle rachis 1 (*Btr1*) (Pourkheirandish et al., 2018) and was supported by the BLAST hit of a coding sequence (Supplementary text S1) of *Btr1* on the reference genome used to genotype our population (*T*. *urartu* pseudomolecule), which was occurred at 62 Mb on chromosome 3A. We also observed that the raw F_ST_ values for three consecutive sites of the region (62 Mb) had the highest (F_ST_=1) values. Thus, this selection scan identified the impact of selection for *Btr1* in the domesticated einkorn.

### A-genome Core Collection

To maximize the utility of the WGRC collection we identified a core set that captured the majority of allelic diversity within 19 *T*. *urartu* accessions, and 60 accessions of *T*. *monococcum* (einkorn wheat) (Supplementary Table S2). In core sets of the entire A-genome collection, we captured ~98 % of the identified alleles, whereas each separate sub-core also captured at least 80% of the segregating alleles of the respective species-specific collections (Table 1). Richness in allelic diversity within the core collections was confirmed by the higher Nei’s diversity index (0.27) of the selected cores relative to the entire collection (0.25) (Table 1). Distribution of the core set accessions in the phylogenetic cluster, PCA clusters, and in the geographic map showed that the selected accessions also represented all subgroups within the population and covered the geographic range (Supplementary Figure S8 – S10). Ranges of glume color scores (Supplementary Table S2) in the core indicated that the core collections are also an excellent representative of phenotypic variations within the whole collection.

## Discussion

### A-Genome Species Distribution, Einkorn Nomenclatures and Morphology

Our results confirm that the WGRC A-genome collection includes arrays of naturally selected germplasm around the center of origin (Figure 1). While verifying the morphologically based grouping of A-genome species through population analysis, we identified three genetically different wild einkorn races (Figure 2 and 3). This information is very crucial in handling a large group of wild einkorn so that accessions with desired genetic background and morphology of interest can be selected for utilization in breeding and further investigation. The wild einkorn genetic races described herein, matched with the races described Kilian et al. (2007), add information to establish the evolutionary and genetic relationships between wild and domesticated einkorn wheats.

However, various nomenclature of the einkorn (Supplementary Figure S11) creates a conundrum in interpreting the different races within the wild einkorn. Some einkorn nomenclature is written in multiple languages; Schiemann (1948) published his nomenclature in German and Dorofeev et al. (1979) in Russian, which could have reduced the acceptance of the nomenclatures by the wider research communities (Dorofeev et al., 1979). In a revised form of MacKey’s classification (Mac Key, 2005b), the *T*. *monococcum* subsp. *boeoticum* was changed to *T*. *monococcum* subsp. *aegilopoides* (Goncharov, 2011). Therefore, no single einkorn classification is deemed to be the most widely accepted and uniformly used. The van Slageren (1994) nomenclature that we follow also is mostly in agreement with the MacKey classification, because both systems use *T*. *monococcum* subsp. *aegilopoides* as the wild einkorn. With all these issues, an updated and widely accepted monograph of einkorn may help maintaining uniformity in taxonomy of these natural accessions.

Species and subspecies classification is first based on morphology. Multiple studies also have discussed different ecogeographical wild einkorn races that have intermediate morphology. Van Zeist (1992) described two groups of wild einkorn: the first group (*T*. *boeoticum* subsp. *thaoudar*) predominately exists in the southeast Turkey, northern Iraq, and west Iran, and the second group (*T*. *boeoticum* subsp. *aegilopoides*) primarily occurs in the west Anatolian center (VAN ZEIST, 1992). The first group of accessions had a double-grained spikelet, and the second group was single-grained, suggesting that the second group is more similar to domesticated einkorn. Brandolini and Heun (2019) explained the *T*. *boeoticum* subsp. *aegilopoides* as an intermediate type feral (semi-wild) einkorn with a semi-brittle rachis and *T*. *boeoticum* subsp. *thaoudar* as the truly wild einkorn with an extremely brittle rachis and argued on the quantitative nature of brittleness in einkorn wheat (Brandolini & Heun, 2019). They hypothesized that the feral einkorn had evolved when agriculture moved from the southeast to west Turkey and Balkans. The semi-brittle rachis breaks into two parts only after being bent and the naturally emerged semi-brittle rachis einkorn mutant still exists in the vicinity of the Karacadag, however, the area is predominant for the truly wild double-grained einkorn (Brandolini & Heun, 2019). Some einkorn accessions with intermediate leaf hairiness, a trait used to classify accessions that is common only in wild einkorn, was also observed (Empilli et al., 2000), indicating that einkorn with intermediate or intergraded morphological characteristics are common (Harlan & Zohary, 1966). The three genetic races of wild einkorn observed in this study also possess unique genetic relationships with cultivated einkorn, as shown by phylogenetic grouping and pairwise F_ST_ values, showing the varying levels of relatedness within and between wild einkorn accession.

### Gene Bank Curation

Globally, plant gene banks often suffer from identified and unidentified duplicates that unnecessarily increase maintenance costs (Díez et al., 2018). Here, we curated 930 A-genome species accessions in WGRC gene bank, identifying duplicates and misclassified accessions, and recognizing valuable unique accessions using genotyping. The existence of misclassified accessions in the gene bank may be due to human error on class assignment and/or data recording; rarely, some accessions might also have controversial morphology. As an example of severe misclassification, consider the wild einkorn accession PI 427328 discussed earlier. Except for the WGRC and Leibniz gene banks, other three gene collection agencies have listed this accession (PI 427328) as *T*. *urartu* (https://www.genesys-pgr.org/a/v2JRrMq2g22), illustrating the importance of genetic scrutiny of the misclassified accessions within the A-genome accessions in different repositories. This genotype-based curation reduces the gene bank operation costs and makes germplasms preservation and utilization easier.

### *T*. *urartu*: the Closest A-genome Diploid Relative of Wheat

With GBS information here we showed that *T*. *urartu* is the closest diploid A-genome relative of wheat and thereby most likely donor of A-genome to the hexaploid wheat (Supplementary Figure S2). This study endorses the known relationships between wheat and A-genome diploids, which was based on thousands of molecular markers and samples (~1,000 diploids and > 200 wheat). Most previous studies describing the relationship between *T*. *urartu* and wheat (Dvořák et al., 1993) however relied on cytogenetic analysis.

### Wild Einkorn Races

The wild einkorn groups were previously divided into α, β, and γ races (Kilian et al., 2007; Zaharieva & Monneveux, 2014)which was consistent with the phylogeny observed in our study. Furthermore, we validated these race groups to match accessions with common USDA PI numbers in both studies overlapped their point of collection and almost all fall under the same races in both studies (Supplementary Table S4). Comparing between studies, there were a few discrepancies in race assignment of accessions (Kilian et al., 2007) that needed correction. For example, Killian et al. (2007) grouped PI 427328 in *T*. *urartu*, but our genetic analysis grouped it into α race within subsp. *aegilopoides* which is also in harmony with WGRC database (accession no. TA879). According to the Genesys database (https://www.genesys-pgr.org/10.25642/IPK/GBIS/98704), another gene bank (Leibniz Institute of Plant Genetics and Crop Plant Research) also classified this PI 427328 within wild einkorn but under the name *T*. *baeoticum* Boiss. subsp. *boeoticum* exemplifying multiple wild einkorn nomenclatures use and creating confusion when describing wild einkorn. Interestingly, Kilian et al. (2007) reported a few feral types of einkorn accessions, indicating they are *T*. *monococcum subsp*. *aegilopoides* according to the nomenclature used, which we did not observe in the WGRC collection. We show that the wild and domesticated einkorn can clearly be differentiated based on genomic data into α, β, and γ races and the domesticated accessions. Given the difficulty and ambiguity of morphological classification, the genetic classification from genomic data can be a preferred approach to cleanly classify any given accession.

### Population Analysis and Different Groups Under A-genome Species

The population structure and F_ST_ analysis on the A-genome species endorsed the established relationships between the species and subspecies. For instance, hybrids between *T*. *monococcum* and *T*. *urartu* are largely sterile and, hence, the genetic differentiation between these species is apparent (Fricano et al., 2014). Also, the intraspecific population differentiation between groups under einkorn at relatively higher K values supported the known genetic relationship between these crossable subspecies that produce mostly fertile hybrids (Harlan & Zohary, 1966).

Our analysis shows that the α race einkorn accessions most likely represent the truly wild einkorn with an extremely brittle rachis, most likely the group of accessions that were traditionally classified as *T*. *boeoticum* subsp. *thaoudar* (Brandolini & Heun, 2019). Differentiation of subpopulations within the α race wild einkorn corresponding to geographic distribution implies migration and genetic drift among truly wild einkorn in the Near East. The *T*. *urartu* subgrouping of accessions from Lebanon and Turkey agrees with Wang et al. (2017), where two subgroups, Mediterranean coastal and Mesopotamia-Transcaucasia, within *T*. *urartu* were reported (Wang et al., 2017).

### Diversity Analysis

Cultivated einkorn had a lower Nei’s diversity index (0.058) than the wild sister group and wild *T*. *urartu* (Table 1), which was expected. As a domesticated species, subsp. *monococcum* experienced a strong population bottleneck and artificial selection might have triggered genetic erosion. On the other hand, the population structure of cultivated einkorn did not show substantial admixture, with the exception of a few accessions, all individuals were true to the ancestry (Figure 2), suggesting a low post domestication admixture contributing elevated diversity. The involvement of a single race (β) in domestication would have further reduced allelic diversity in the cultivated einkorn; there was no difference between the Nei’s diversity of β race (0.058) and the domesticated einkorn (0.058). Kilian et al. (2007) illustrated no nucleotide diversity was reduced during einnkorn domestication; instead, they observed increased diversity in domesticated compared to wild einkorn (Kilian et al., 2007). However, the diversity assessment in (Kilian et al., 2007) could be influenced by the limited number of loci and smaller sample size; especially, diversity estimates are sensitive to sample size when there are only a handful of markers (Bashalkhanov et al., 2009; Li et al., 2009). In this experiment, we used thousands of SNP markers and have larger sample size, which minimized the effect of sample size and the number of loci. The highest Nei’s diversity index (0.25) for all A-genome combinedly, and the considerably higher Nei’s diversity index for each species and core collections indicated that these accessions are very important assets with novel and useful genetic variations.

### *Btr1*: Einkorn Domestication Signal

Through F_ST_ computation, we showed that in einkorn wheat there is a single strong selection signal observed on chromosome 3A corresponding to the *Btr1* locus (Supplementary Figure S7). Previous study also described *Btr1* as one of the most important features of einkorn domestication (Pourkheirandish et al., 2018). The non-brittleness in domesticated einkorn is controlled by a single nucleotide change in *Btr1* of wild einkorn that results in an amino acid substitution (alanine to threonine) (Pourkheirandish et al., 2018). With ~ 1,000 filtered loci per chromosome, we located the candidate selection region. The availability of a *T*. *monococcum* reference genome to call the genotype would be ideal for obtaining dense markers and better locating the selection signature on einkorn wheat.

### Core Collections

Establishing core collections of A-genome species enabled harnessing useful genetic variation to improve wheat and cultivated einkorn. To the best of our knowledge, this is the first genetic core of A-genome species, which included only 79 accessions and yet contains ~ 98% of the identified alleles while achieving a more than 10-fold (79/930) reduction in the number accessions (Table 1, Supplementary Table S2). The Nei’s diversity index computed for these core collections supported that they have considerably higher relative diversity and can be leveraged for targeted germplasm improvement.

## Conclusions

This study reports the important aspects of the A-genome wheat species for genetic diversity, gene bank curation, and core set selection. Following an assessment of nearly 1,000 accessions, we report that the A-genome species possess a considerable amount of genetic diversity, which can be utilized in breeding wheat and domesticated einkorn. This vast diversity is most effectively managed in pre-breeding with well-defined core collections. Identifying and in-depth characterizing of such core collections adds significant value and accessibility to the germplasm. Having a well curated and accurately described gene bank collection, as done here, is a critical foundation to effectively using this rich diversity for crop improvement and enhancing the value of gene bank resources.

## Author Contributions

JP and JR designed the experiment and conceptualized the study. JR, SW, DW, BE and NS carried out experiments and data collection, growing of plant materials and germplasm. LA conducted data analysis. LA and JP wrote the manuscript. All the authors have read and approved the manuscript for publication.

## Funding information

This material is based upon work supported by the National Science Foundation and Industry Partners under Award No. (1822162) “Phase II IUCRC at Kansas State University Center for Wheat Genetic Resources” and the National Science Foundation under Grant No. (1339389) “GPF-PG: Genome Structure and Diversity of Wheat and Its Wild Relatives”. Any opinions, findings, and conclusions or recommendations expressed in this material are those of the author(s) and do not necessarily reflect the views of the National Science Foundation or industry partners.

## Materials and Methods

### Plant Resources

This study included 930 accessions of the A-genome diploid wheat species maintained in the WGRC gene bank (Supplemental Table S1), which were primarily acquired from the Near East, Transcaucasia, and the Balkans (Figure 1). Most of the A-genome accessions (~ 85%) tested include those initially collected by B. Lennert Johnson, University of California–Riverside in the 1960s and 1970s. The remaining accessions were obtained from gene banks in Japan (22), Germany (24), and ICARDA (61). Several accessions were donated by Robert Metzger, USDA, Oregon State University, Corvallis (26), seven were collected by the WGRC, and the remainder (10) from other sources. We also tested 225 CIMMYT wheat lines (Supplementary Table S1) genotyped earlier with GBS SNPs (Gao et al., 2021) thereby inferring the genetic relationships between A-genome diploids and the wheat.

### Genotypic Characterization

The tested accessions were grown as single plants in the greenhouse and tissue collected in 96-well plates. The tissues were lyophilized for ~3 days and ground to a fine powder using Retsch mixer mill MM 400. Genomic DNA extraction and GBS library preparation steps were according to Singh et al. (2019). We had total four multiplexed GBS libraries including one for the pilot study. The pilot study GBS library was 384-plex, whereas the other GBS libraries were 288-plex. We sequenced on the Illumina platform with 150 bp pair-end reads (PE150). We had total The information about GBS of 226 CIMMYT lines can be obtained (Gao et al., 2021).

The TASSEL5 GBSv2 pipeline was used for sequence data processing and genotype calling (Glaubitz et al., 2014). Reads were aligned to a *T*. *urartu* pseudomolecule reference (Ling et al., 2018) using bowtie2 alignment (Ling et al., 2018) and exported to variant call format (VCF). Filtering of the VCF was done for bi-allelic SNPs using the Fisher exact test with a threshold p-value <0.001 as described previously (Poland et al., 2012b) considering that the genotypes should represent biallelic variants in inbred accessions. Genotypes for accessions across all A-genome species were called together, followed by extracting variants segregating within each species using VCFtools (Danecek et al., 2011). The SNPs were filtered for minor allele frequency (MAF) > 0.01, missing percentage < 30%, and heterozygous genotypes < 10% at the population level using TASSEL and R (R Core Team 2019).

The A-genome diploids and wheat lines were genotyped together calling SNP on A-genome of wheat reference genome of cultivar Chinese Spring (iwgsc_refseqv1.0) (Appels et al., 2018). We also filtered these SNPs using aforementioned criteria. The unrooted neighbor-joining (NJ) phylogenetic tree of A-genome diploid and wheat lines were generated for investigating the genetic relationship. We followed approach of Singh *et al*. (2019) to generate NJ tree from GBS sampled population, where clustering was conducted with default parameters of R packages ‘dist’, ‘ape’, and ‘phyclust’.

### Gene Bank Curation

A-genome species in the WGRC gene bank were curated to identify misclassified and duplicate accessions. The misclassified accessions identified based on the genetic properties were compared with accessions in the adjusted class morphologically to assure if they were previously assigned or documented to the wrong class. Furthermore, to confirm the ploidy of the misclassified accessions that were grouped far from the major *T*. *urartu* clade and did not exhibit a closer relationship with any diploid A-genome in genetic tree, chromosome counts were made by staining with 4’,6-diamidino-2-phenylindole (DAPI). The detail method for chromosome count was obtained (Koo et al., 2017).

The genetically identical accessions were identified using pairwise allele matching across homozygous and non-missing sites. We first analyzed the loci identity proportions distribution at genome-wide scale including every possible pair-wise comparison among accessions within a single species. A threshold for allele matching percentage given discrepancies for sequencing errors was then detected by finding a point that separates the local maxima existing around the prefect identity (100%). The identity matrix and percentage allele matching were computed in R using a custom script as described by (Singh et al., 2019). The morphological similarity and the geographical relations of the identified duplicate accessions were checked for confirmation. Glume color (level of darkness) was used as a morphological marker for cross-validation to affirm the accessions in a duplicate set have the same or similar phenotypes. The variation in glume color was rated from completely white (0) to dark black (9).

### Population Structure

Population structure of A-genome wheat species was analyzed using fastStructure (Raj et al., 2014). The fastStructure was initially run at K=2 to K=12 with three replications using ‘simple’ prior where K refers to number of population or model complexity. For the optimum value of K, the program was run using ‘logistic’ prior at K=2 to K=7 with three replications (Singh et al., 2019). An appropriate number of K was also obtained using the fastStructure provided utility tool, chooseK.py. The fastStructure output was graphically visualized using an R package POPHELPER (Francis, 2017). Passport information including the classification based on morphology, and the accessions geographical sites were used to group and reorder the samples in population analysis. Accessions that were identified as misclassified were confirmed through morphological evaluation and reordered to subspecies based on the genotype-based grouping and the final result was plotted.

Phylogenetic clustering was carried out in R using ‘dist’ function and ‘ape’ and ‘phyclust’ packages (Singh et al., 2019). The branches of an unrooted neighbor-joining (NJ) tree were first colored using the morphology-based classification, and then according to genotype analysis. The morphology-based coloring was particularly focused in identifying misclassified accessions. A-genome species population genetic structure was also dissected using principal component analysis (PCA) of genomic data. For PCA, we estimated the eigenvalues and eigenvectors on R using the ‘e’ function in ‘A’ matrix obtained from the rrBLUP (Endelman, 2011; Singh et al., 2019).

### Analysis of Genetic Diversity

A-genome species genetic diversity was assessed by computing the Nei’s diversity index (Nei 1973) using filtered genotyping markers (Nei, 1973). We computed the Nei’s indices of (1) all A-genome accessions together, (2) each species and subspecies independently, (3) the races within the subspecies, and (4) and the core collections. The minor allele frequency (MAF) for each species was also plotted to discern the excess of rare variants in respective population. Number of segregating loci per group were determined (Table 1). A pairwise fixation index (F_ST_) (Nei, 1987) also was computed between the species and subgroups separated by the population analysis (Singh et al., 2019).

### F_ST_ Within Einkorn and Selection signature

We computed a genome-wide F_ST_ statistic for variants within the einkorn group using R (R Core Team 2019) as described (Porto-Neto et al., 2013). This method compute F_ST_ statistic based on pure drift model (Nicholson et al., 2002). We also compared the output by computing the Cockerham and Weir F_ST_ statistic (Weir & Cockerham, 1984) using VCFtools (Danecek et al., 2011). The *T*. *monococcum* VCF file with biallelic variant was further filtered keeping SNPs with MAF > 0.01, missing < 30% and heterozygous < 10% followed by imputation using Beagle 5.1 (Browning et al., 2018). The filtered and imputed genotyping information was used to derive the F_ST_ values. To balance the population sizes of domesticated and wild einkorn, we randomly chose 145 wild einkorn accessions to match the number of 145 domesticated accessions. The F_ST_ were plotted using ggplot2 in R (R Core Team 2019) and the raw F_ST_ plots were smoothed using Lowess method (Pintus et al., 2014) to find the genomic regions with extreme F_ST_. To define the selection signal peak, we considered outlier F_ST_ values that were more than three standard deviation (3σ) over genome wide average as the threshold.

### Core Collections

Core collections of *T*. *urartu*, and *T*. *monococcum* (wild and domesticated einkorn) were selected taking allelic diversity, genotype coverage, geographical representation, and phenotypic variation (glume color) into consideration. From the filtered genotyping file, heterozygous genotypes were masked before running the core accessions selection software GenoCore (Jeong et al., 2017). We ran GenoCore with the default parameters: -d 0.01% and -cv 99%. The positions of the selected samples within the phylogenetic tree and PCA clusters were observed through coloring the selected core accessions versus all other samples. Also, the geographical representations were evaluated marking the selected vs. remaining accessions in the google map using GPS Visualizer (https://www.gpsvisualizer.com). To ensure phenotypic variations in the selected core sets, we considered the glume color score (Supplementary Table S2) as a reference variation. The Nei’s diversity index (1987) of core sets were also computed (Danecek et al., 2011).

## Accession Numbers

Raw sequence data obtained from GBS, the fastq files, has been deposited at the National Center for Biotechnology Information (NCBI) SRA database with the BioProject accession PRJNA744683 (https://www.ncbi.nlm.nih.gov/sra/PRJNA744683). The GBS key file with required information for demultiplexing and further detail about the SRA deposited fastq files can be obtained at Dryad digital repository (doi:10.5061/dryad.9zw3r22f6).

## Supplemental Data

**Supplementary Table S1**. List of A-genome accessions, their origin and duplicated accessions.

**Supplementary Table S2**. Core collections of A-genome species.

**Supplementary Table S3**. The misclassified A-genome species accessions, their previous class based on morphology and the updated class/group based on the genotyping.

**Supplementary Table S4**. Number of accessions with common PI numbers that clustered in corresponding groups in this experiment and a past experiment. Both studies tested only a portion of germplasms from USDA. The α, β, and γ races indicate the three genetic clusters within the wild einkorn as designated in the past study. The * indicates the accessions that we detected as misclassified and need adjustment of class. Past study also grouped these accessions in the same group that we observed, however, they did not discuss on misclassification issue and just listed the accessions based on morphological classification.

**Supplementary Table S5**. Pairwise F_ST_ coefficients among the subgroups within α race of subsp. *aegilopoides* (wild einkorn) and the admixture groups. There were three subgroups (Turkey, Duhok, Erbil) and two admixture groups (Iran and Sulaymaniyah (Iraq)).

**Supplementary Figure S1**. The *T*. *urartu* clade and subsp. *aegilopoides* α race clade in the unrooted NJ tree highlighting the misclassified accessions between the two groups. The red branches within the gold-colored clade and the gold branches within the red clade reflect the misclassified accessions.

**Supplementary Figure S2**. Threshold determination for declaring duplicate accessions identification. (A) Percentage identity versus the number of comparisons among 204 accessions in *T*. *urartu* (D) Percentage identity versus the number of pairwise comparisons for the accession pairs in *T*. *urartu* that had near perfect (≥ 99%) identity.

**Supplementary Figure S3**. An unrooted Neighbor-Joining (NJ) tree of wheat and A-genome species: *T*. *urartu*, subsp. *aegilopoides*, and subsp. *monococcum*. The tree branches are colored based on the genetic grouping of the accessions after correcting misclassified accessions. *T*. *urartu* (yellow), domesticated einkorn (red), wild einkorn race α (blue), and wild einkorn race γ (green), and misclassified tetraploids (brown) are shown.

**Supplementary Figure S4**. The mitotic metaphase cell of a misclassified wild wheat accession in WGRC collection, TA10881 confirming the accession as tetraploid (2n=4x=28) and thus verified the GBS based genetic grouping. Chromosomes were stained with 4’,6-diamidino-2-phenylindole (DAPI). Before genotyping and cytological confirmation, the accession was falsely grouped under *T*. *urartu*.

**Supplementary Figure S5**. Principle component analysis (PCA) plot for A-genome wheat species with two major PCs. There were three races α, γ and β within the wild einkorn group which clustered separately in population analysis.

**Supplementary Figure S6**. Minor allele frequency plots of A-genome diploid species: (**a**) *T*. *monococcum* subsp. a*egilopoides*, (**b**) *T*. *monococcum* subsp. *monococcum*, and (**c**) *T*. *urartu*

**Supplementary Figure S7**. Smoothed F_ST_ curve showing selection signal for einkorn wheat on chromosome 3A. The strongest signal was located at 60-90 Mb. The horizontal green line indicates the selection signature determination genome-wide threshold (0.24), which is 3σ above the mean. The red vertical line at 62 Mb on chromosome 3A indicates the location of candidate selection signature Btr1.

**Supplementary Figure S8**. An unrooted NJ phylogenetic tree of A-genome wheat species showing the accessions in the core collections and all other accessions in respective clades. Black branch reflects the accessions in the core collection, and the golden branch indicates all other accessions that are not in the core collections.

**Supplementary Figure S9**. Principle component analysis (PCA) plot of A-genome wheat accessions showing partitioning of different groups within the species and the accessions selected in the genetic cores (black triangles).

**Supplementary Figure S10**. Geographic map of A-genome wheat accessions, where the core accessions were indicated by larger google marks and the rest of the accessions were shown by smaller marks of the respective groups.

**Supplementary Figure S11**. Diagram showing three different taxonomic classification systems of einkorn wheat. In WGRC, we follow the taxonomic classification system of Van Slageren (1994).

**Supplementary text S1**. Coding sequence of gene for non-brittle rachis 1 (Btr1) in *T*. *monococcum* subsp. *monococcum*

## Acknowledgements

We would like to thank the wheat genetic resource center (WGRC) gene bank for collecting and maintaining these A-genome species.

Authors have no competing interests

All data are available in the manuscript or the supplementary resources

